# Gnotobiotic rainbow trout (*Oncorhynchus mykiss*) model reveals endogenous bacteria that protect against *Flavobacterium columnare* infection

**DOI:** 10.1101/2020.06.19.161471

**Authors:** David Pérez-Pascual, Sol Vendrell-Fernández, Bianca Audrain, Joaquín Bernal-Bayard, Rafael Patiño-Navarrete, Vincent Petit, Dimitri Rigaudeau, Jean-Marc Ghigo

## Abstract

The health and environmental risks associated with antibiotic use in aquaculture have promoted bacterial probiotics as an alternative approach to control fish infections in vulnerable larval and juvenile stages. However, evidence-based identification of probiotics is often hindered by the complexity of bacteria-host interactions and host variability in microbiologically uncontrolled conditions. While these difficulties can be partially resolved using gnotobiotic models harboring no or reduced microbiota, most host-microbe interaction studies are carried out in animal models with little relevance for fish farming. Here we studied host-microbiota-pathogen interactions in a germ-free and gnotobiotic model of rainbow trout (*Oncorhynchus mykiss*), one of the most widely cultured salmonids. We demonstrated that germ-free larvae raised in sterile conditions displayed no significant difference in growth after 35 days compared to conventionally-raised larvae, but were extremely sensitive to infection by *Flavobacterium columnare*, a common freshwater fish pathogen causing major economic losses worldwide. Furthermore, re-conventionalization with 11 culturable species from the conventional trout microbiota conferred resistance to *F. columnare* infection. Using mono-re-conventionalized germ-free trout, we identified that this protection is determined by a commensal *Flavobacterium* strain displaying antibacterial activity against *F. columnare*. Finally, we demonstrated that use of gnotobiotic trout is a suitable approach for the systematic identification of both endogenous and exogenous probiotic bacterial strains that may protect teleostean hosts against *F. columnare* and other pathogens. This study establishes a novel and ecologically-relevant gnotobiotic model that will improve the sustainability and health of aquaculture.

## INTRODUCTION

As wild fish stock harvests have reached biologically unsustainable limits, aquaculture has grown to provide over half of all fish consumed worldwide [1]. However, intensive aquaculture facilities are prone to disease outbreaks and the high mortality rate in immunologically immature juveniles, in which vaccination is unpractical, constitutes a primary bottleneck for fish production [2-4]. These recurrent complications prompt the prophylactic or therapeutic use of antibiotics and chemical disinfectants to prevent fish diseases [5, 6] but may lead to final consumer safety risks, environmental pollution and spread of antibiotic resistance [7]. In this context, the use of bacterial probiotics to improve fish health and protect disease-susceptible juveniles is an economic and ecological sensible alternative to antibiotic treatments [8, 9].Probiotics are live microorganisms conferring health benefits on the host via promotion of growth, immuno-stimulation or direct inhibition of pathogenic microorganisms [10, 11]. The native host microbiota plays a protective role against pathogenic microorganisms by a process known as colonization resistance [12, 13]. In fish, the endogenous microbial community, whether residing in gastrointestinal tract or in the fish mucus, was early considered as a source of protective bacteria [14-18]. However, selection of probiotic bacteria is often empirical or hampered by the poor reproducibility of *in vivo* challenges, frequently performed in relatively uncontrolled conditions with high inter-individual microbial compositions [15, 19].To improve evidence-based identification of fish probiotics and their efficacy in disease prevention, the use of germ-free (GF) or fully controlled gnotobiotic hosts is a promising strategy [20, 21]. In addition to laboratory fish models such as zebrafish (*Danio rerio*) [22-24], several fish species have been successfully reared under sterile conditions to test probiotic-based protection against pathogenic bacteria, including Atlantic cod (*Gadus morhua*) [25], Atlantic halibut (*Hippoglossus hippoglossus*) [26], European sea bass (*Dicentrarchus labrax*) [19] and turbot (*Scophthalmus maximus*) [27] (for a review, see [28]).

Salmonids, especially rainbow trout (*Oncorhynchus mykiss*) and Atlantic salmon (*Salmo salar*), are economically important species, whose production in intensive farming is associated with increased susceptibility to diseases caused by viruses, bacteria and parasites [29]. Here we studied the probiotic potential of endogenous members of the rainbow trout microbiota to protect against infection by *Flavobacterium columnare*, a fresh-water fish pathogen causing major losses in aquaculture of fish such as Channel catfish, Nile tilapia and salmonids [30]. We developed a new protocol to rear GF trout larvae and showed that GF larvae were extremely sensitive to infection by *F. columnare*. We then identified two bacterial species originating either from the trout microbiota (a commensal *Flavobacterium* sp.) or the zebrafish microbiota (*Chryseobacterium massiliae*) that fully restored protection against *F. columnare* infection. Our *in vivo* approach opens perspectives for the rational and high throughput identification of probiotic bacteria protecting rainbow trout and other fish against columnaris disease. It also provides a new model for the study of host-pathogen interactions and colonization resistance in a relevant teleostean fish model.

## RESULTS

### Germ-free trout show normal development and growth compared to conventional larvae

To produce microbiologically controlled rainbow trout and investigate the potential protection conferred by endogenous or exogenous bacteria against incoming pathogens, we produced (GF) trout larvae by sterilizing the chorion of fertilized eggs with a cocktail of antibiotics and antifungals, 0.005 % bleach and a iodophor disinfection solution. GF eggs were then kept at 16°C under sterile conditions and both conventional (Conv) and treated eggs hatched spontaneously 5 to 7 days after reception, indicating that the sterilization protocol did not affect the viability of the eggs. However, hatching efficiency was 72 ± 5.54 % for sterilized eggs versus 48.6 ± 6.2 % for non-treated, Conv eggs, possibly due to higher susceptibility of Conv eggs to opportunistic infections from the endogenous microbiota. Once hatched, all larvae were transferred into vented-cap cell culture flasks containing fresh sterile water without antibiotics renewed every 48 hours (h). GF and Conv fish relied on their vitellus reserves until day 20 days post-hatching (dph) after which they were fed with sterilized fish food powder every 48 h (Fig. 1). Sterility tests were performed at 24 h, 7 days and 21 days post-sterilization treatment and before each water change until the end of the experiment (35 dph) (Supporting Fig. S1). To test the physiological consequences of raising GF larvae, we compared the growth of Conv and GF larvae reared from the same batch of fertilized eggs and observed no significant difference in standard body length (2.51 ± 0.24 cm *vs*. 2.58 ± 0.21 cm) or weight (1.17 ± 0.20 g *vs*. 1.17 ± 0.10 g) at 35 dph for Conv and GF, respectively (Supporting Fig. S2). To compare Conv and GF trout anatomy, we developed an approach combining iDISCO solvent-based method to generate transparent fish tissue and lightsheet 3D imaging of the whole body. This analysis did not reveal any anatomical differences at 21 dph, even regarding organs in direct contact with fish microbiota such as gills (Fig. 2D and 2I) and intestine (Fig. 2C and 2H; Supporting Fig. S3). No difference was seen on other organs potentially influenced by gut-microbiota such as the brain (Fig. 2A and 2F), spleen (Fig. 2B and 2G) and head kidney (Fig. 2E and 2J) [31]. These results suggested that the natural microbiota had no major macroscopic impact on fish growth, development or anatomy at this stage of rainbow trout development in our rearing conditions.

**Figure 1.**
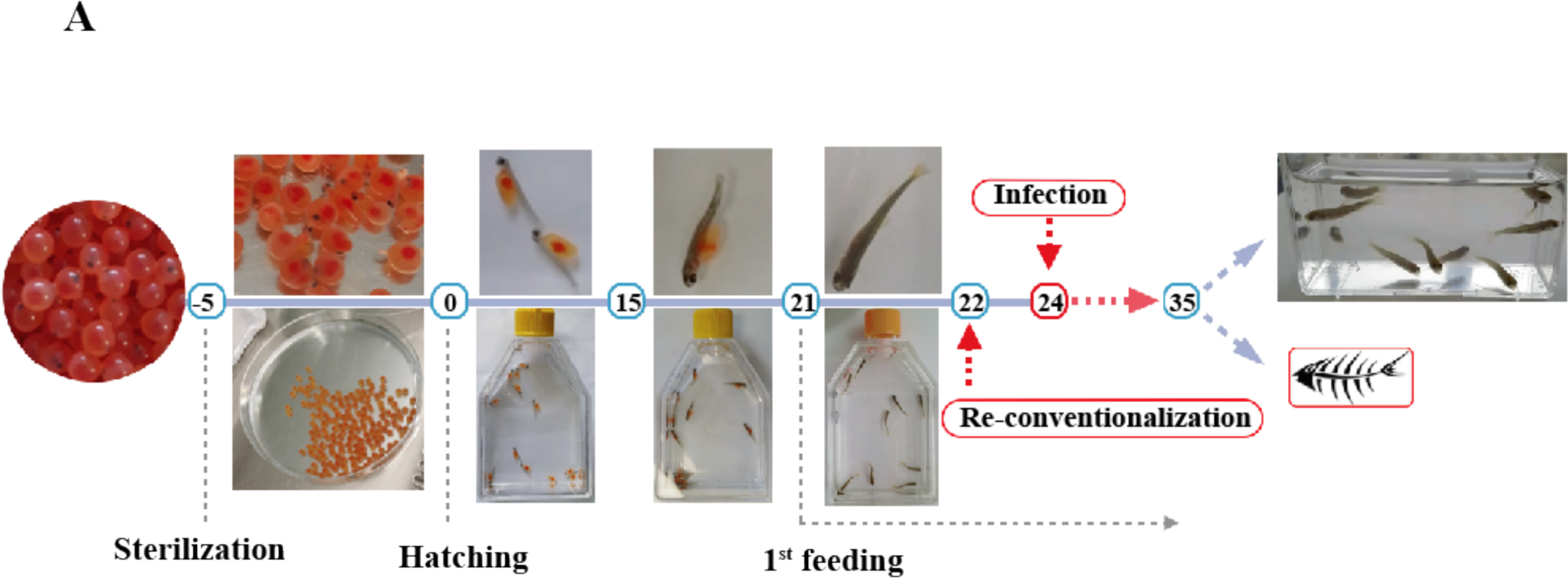
Protocol used in this study to raise and infect or re-conventionalize germ-free (GF) trout larvae. Eyed eggs were sterilized 5 days before hatching (−5 dph) and kept in sterile, autoclaved mineral water at 16°C in Petri dishes until hatching. Once hatched, rainbow trout larvae were transferred into vented cap cell culture flasks for the duration of the experiment. Larvae were fed every 2 days with sterile powder food from 21 dph until the end of the experiment; water was renewed 30 minutes after feeding. To test the protective effect of potential probiotic strains, larvae were re-conventionalized by one or several commensal bacteria diluted in water at 22 dph. Pathogenic bacteria were added to the water at 24 dph for 24 h and then larvae were washed with fresh sterile water. Survival after infection was monitored twice per day.

**Figure 2.**
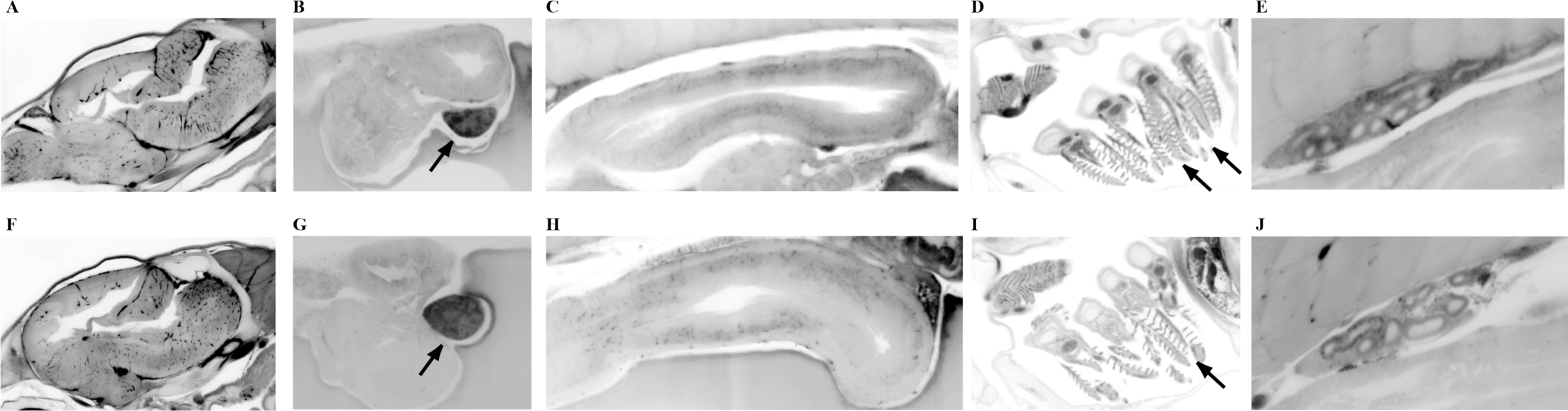
Anatomical comparison of Conventional (Conv) and GF rainbow trout larvae. 3D deep imaging of whole trout body corresponding to autofluorescence signal acquired by lightsheet microscopy after novel fish clearing processing. Selected optical sections of 21 dph were presented for Conv (A, B, C D and E) and GF (F, G, H, I and J) rainbow trout larvae. Brain (A and F), spleen (black arrow in B and G), gut (C and H) (see also supplementary figure S3), gills (black arrows in D and I), and head kidney (E and J). Images representative of two different fish per condition.

### Identification of susceptibility to fish pathogens in germ-free but not conventional trout larvae

To identify bacterial pathogens able to infect GF rainbow trout larvae by the natural infection route, we exposed the 24 dph larvae for 24 h to 10^7^ colony forming units (CFU)/ml of several trout bacterial pathogens, including *Flavobacterium psychrophilum* strain THCO2-90, *F. columnare* strain Fc7, *Lactococcus garvieae* JIP 28/99, *Vibrio anguillarum* strain 1669 and *Yersinia ruckeri* strain JIP 27/88 [32]. Larvae were then washed with sterile water, renewing 90% of the infection water three times and kept at 16°C under sterile conditions. Among all tested pathogens, only *F. columnare* strain Fc7 led to high and reproducible mortality of GF trout larvae within 48 h post-exposure (Fig. 3). In contrast, Conv larvae reared from non-sterilized eggs survived *F. columnare* strain Fc7 infection under tested conditions (Fig. 4A). Histological analysis performed at 25 dph (24 h post infection) on GF and Conv larvae did not show any sign of intestinal damage (Supporting Fig.S4).

**Figure 3.**
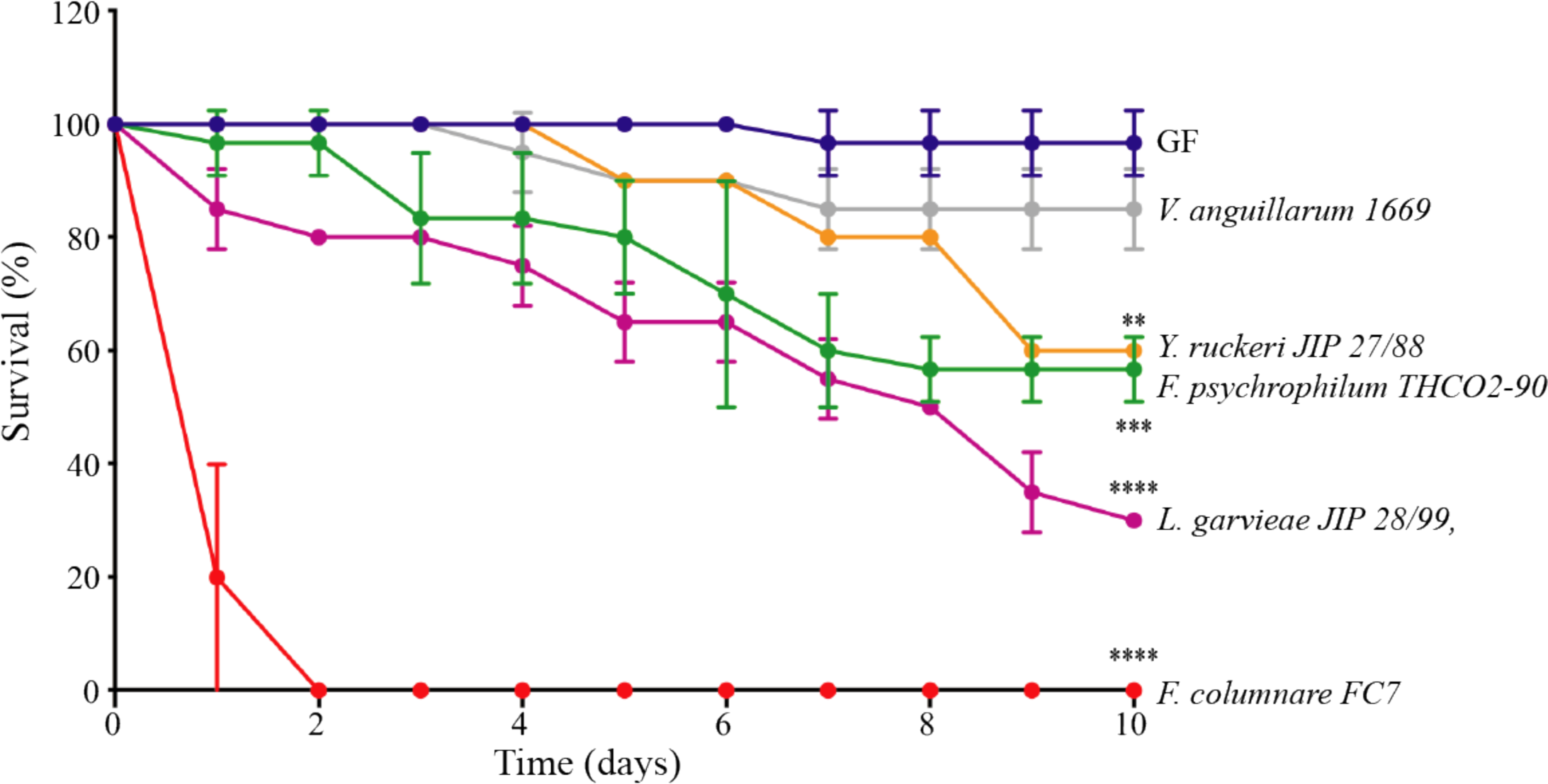
Survival of GF and Conv rainbow trout larvae infected with different fish pathogens. Kaplan-Meier graph of GF larvae survival after bath exposure to *F. psychrophilum* strain THCO2-90, *F. columnare* strain Fc7, *L. garvieae* strain JIP 28/99, *V. anguillarum* strain 1669 and *Y. ruckeri* strain JIP 27/88. Mean and SD plot representing average survival percentage of fish for 10 days after exposition to different pathogenic microorganisms. For each condition n = 10 larvae. All surviving fish were euthanized at day 10 post-infection. Asterisks indicate significant difference from non-infected population (**p<0.01; ***p<0.001; ****p<0.0001).

**Figure 4.**
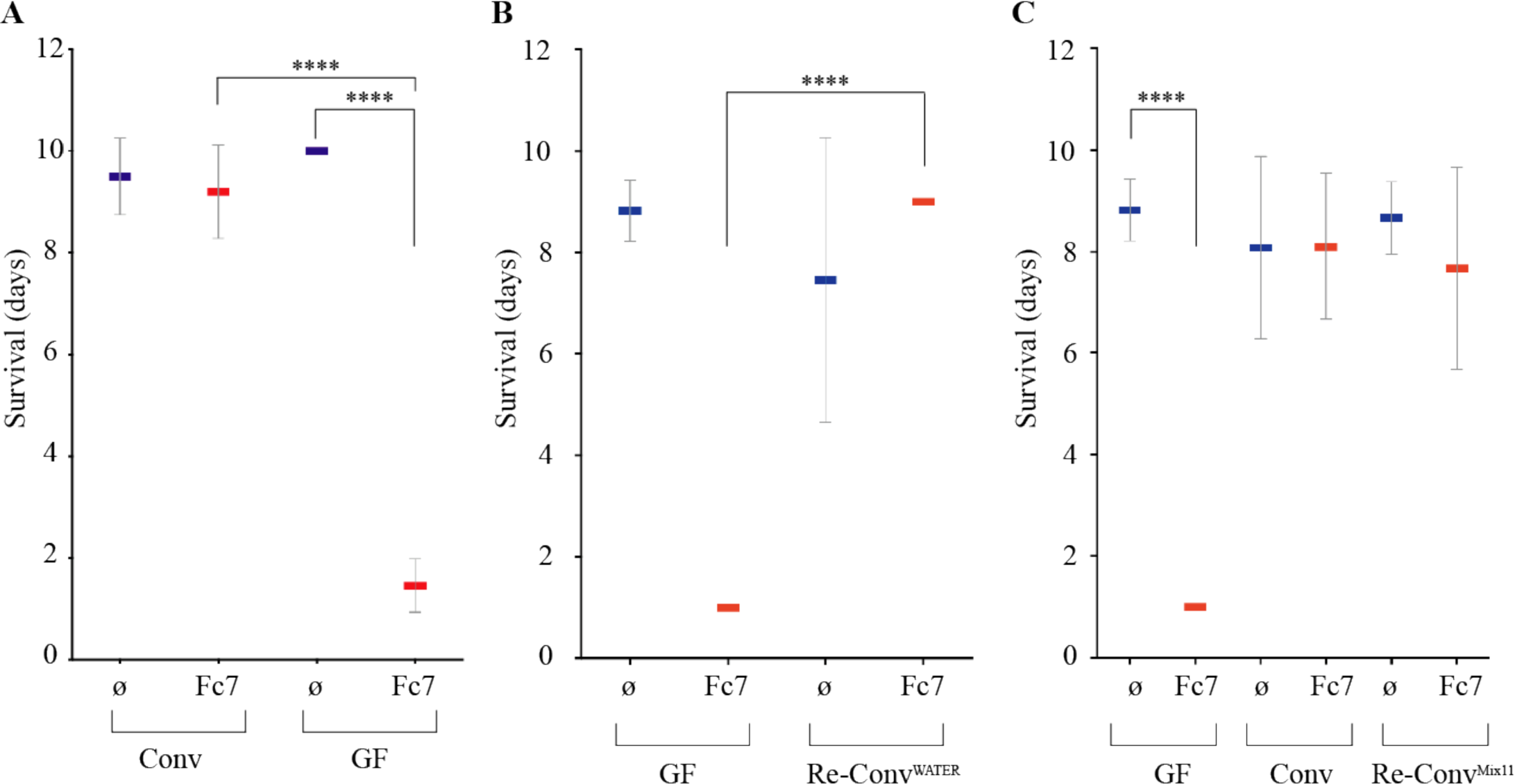
Survival of re-conventionalized trout larvae against *F. columnare* Fc7 infection. **A:** *F. columnare* strain Fc7 kills GF but not Conv rainbow trout. Mean and SD plot representing average day post-infection at which infected fish die. For each condition n = 10 larvae. All surviving fish were euthanized at day 10 post-infection. Asterisks indicate significant difference from non-infected population (****p<0.0001). **B:** GF trout larvae exposed to water used to raise Conv fish at 21 dph show similar survival rates to *F. columnare* infection compared to Conv trout larvae. **C:** The 11 strains identified from Conv fish microbiota were added to rainbow trout larvae at 22 dph, followed by *F. columnare* infection at 24 dph. This bacterial mixture protected re-conventionalized larvae from infection. For each condition n = 10 larvae. All surviving fish were euthanized at day 10 after infection (****p<0.0001).

### Conventional rainbow trout microbiota protects against *F. columnare* infection

Considering the high sensitivity of GF but not Conv trout larvae to infection by *F. columnare* Fc7, we hypothesized that resistance to infection could be provided by some components of the Conv larvae microbiota. To test this, we exposed GF rainbow trout larvae to water from Conv larvae flasks at 21 dph. Re-conventionalized (Re-Conv) rainbow trout larvae survived as well as Conv larvae to *F. columnare* Fc7 infection, whereas those maintained in sterile conditions died within the first 48h after infection (Fig. 4B). These results suggested that microbiota associated with Conv rainbow trout provide protection against *F. columnare* Fc7 infection. To identify culturable species potentially involved in this protection, we plated bacteria recovered from 3 whole Conv rainbow trout larvae at 35 dph on various agar media. 16S rRNA-based analysis of each isolated morphotype led to the identification of 11 different bacterial strains corresponding to 9 different species that were isolated and stored individually (Table 1).

**Table 1.** The 11 strains isolated from Conv rainbow trout larvae.

| Bacterial strains isolated from trout microbiota |
| --- |
| *Aeromonas rivipollensis* 1 |
| *Pseudomonas helmanticensis* |
| *Aeromonas rivipollensis* 2 |
| *Pseudomonas baetica* |
| *Aeromonas hydrophila* |
| *Flavobacterium plurextorum* 1 |
| *Acinetobacter* sp. |
| *Flavobacterium plurextorum* 2 |
| *Delftia acidovorans* |
| *Flavobacterium* sp. strain 4466 |
| *Pseudomonas* sp. |

We then re-conventionalized GF rainbow trout larvae at 22 dph with an equiratio mix of all 11 identified bacterial strains (hereafter called Mix11), each at a concentration of 5.10^5^ CFU/ml. After exposure to *F. columnare* strain Fc7, these Re-Conv^Mix11^ larvae survived as well as Conv fish (Fig. 4C), demonstrating that the Mix11 isolated from the rainbow trout microbiota recapitulates full protection against *F. columnare* infection observed in Conv larvae.

### Resistance to *F. columnare* infection is conferred by one member of the trout microbiota

To determine whether some individual members of the protective Mix11 could play key roles in infection resistance, we mono-re-conventionalized 22 dph GF trout by each of the 11 isolated bacterial strains at 5.10^5^ CFU/ml followed by challenge with *F. columnare* Fc7. We found that only *Flavobacterium* sp. strain 4466 restored Conv-level protection, whereas the other 10 strains displayed no protection, whether added individually (Fig. 5A) or as a mix (Mix10 in Fig. 5B). Interestingly, although cell-free spent supernatant of *Flavobacterium* sp. strain 4466 showed no inhibitory activity against *F. columnare* Fc7 in an overlay assay (Supporting Fig. S5A), *Flavobacterium* sp. strain 4466 colony growth inhibited the growth of *F. columnare* Fc7 (Supporting Fig. S5B) and of all tested *F. columnare* strains (Supporting Fig. S5C), suggesting a potential contact dependent inhibition. Consistently, we identified a cluster of 12 genes in the *Flavobacterium* sp. strain 4466 genome (*tssB, tssC, tssD, tssE, tssF, tssG, tssH, tssI, tssK, tssN, tssP* and *tssQ*) characteristic of type 6 secretion system (T6SS), T6SS^iii^, a contact-dependent antagonistic system only present in phylum *Bacteroidetes* [33]. To improve the taxonomic identification of the protective *Flavobacterium* isolated from the trout larvae microbiota, we performed whole genome sequencing followed by Average Nucleotide Identity (ANI) analysis. We determined that despite similarity with *Flavobacterium spartansii* (94.65 %) and *Flavobacterium tructae* (94.62 %), these values are lower than the 95 % ANI needed to identify two organisms as the same species [34]. Furthermore, full-length 16S rRNA and *recA* genes comparisons also showed high similarity with *F. spartansii* and *F. tructae*, however, the obtained values were also below the 99 % similarity threshold required to consider that two organisms belong to the same species (Supporting Table S1). Similarly, a maximum likelihood based phylogenetic tree (Supporting Fig.S6) generated from sequences of 15 bacterial strains from the *Flavobacterium* genus revealed that the sequence of *Flavobacterium* sp. strain 4466 clustered with sequences of *F. spartansii* and *F. tructae*, but did not allow the identification of *Flavobacterium* sp. strain 4466 at species level.

**Figure 5.**
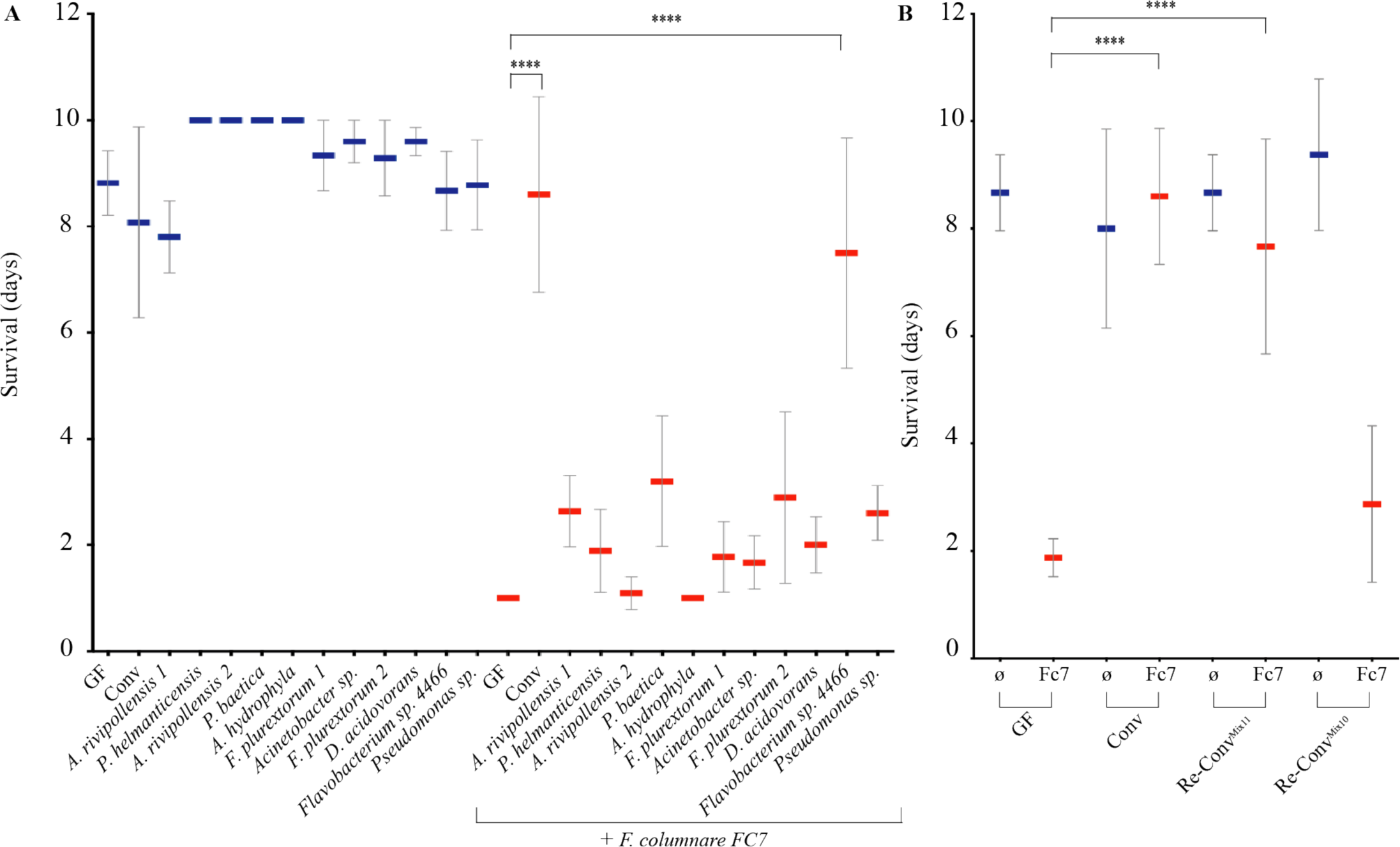
Protection of GF trout larvae against *F. columnare* infection by individual species isolated from the Conv rainbow trout microbiota. **A:** The 11 species isolated from Conv fish microbiota (Table 1) were added individually to rainbow trout larvae at 22 dph, followed by *F. columnare* Fc7 infection at 24 dph. From the 11 different strains, only *Flavobacterium* sp. strain 4466 protected re-conventionalized larvae from infection. **B:** Mix11, Mix10 (mix of all identified strain with the exception of *Flavobacterium* sp. strain 4466), were added to rainbow trout larvae at 22 dph, followed by *F. columnare* infection at 24 dph. Mix11 protected re-conventionalized larvae from infection, whereas Mix10 did not. For each condition n = 10 larvae. All surviving fish were euthanized at day 10 after infection (****p<0.0001).

### Endogenous *Flavobacterium* sp. strain 4466 protects germ-free zebrafish larvae against *F. columnare* infection

*F. columnare* infects a wide range of wild and cultured freshwater fish species [30] and we previously established that GF zebrafish larvae are highly sensitive to *F. columnare* infection [35]. To test whether the protective *Flavobacterium* sp. isolated from the Conv rainbow trout microbiome could also protect zebrafish, we re-conventionalized GF zebrafish larvae with *Flavobacterium* sp. 48 hours before exposure to four virulent *F. columnare* strains (Fc7, ALG-00-530, IA-S-4, and Ms-Fc-4) belonging to genomovars I and II, and isolated from different geographical origins and host fish species. Whereas all tested *F. columnare* strains were highly virulent and killed GF zebrafish larvae within 48 hours, the non-pathogenic *Flavobacterium* sp. strain 4466 conferred protection to all pathogenic *F. columnare* strains except strain Ms-Fc-4 (Figure 6). Therefore, the *Flavobacterium* sp. strain identified from trout Mix11 is a putative probiotic useful beyond trout to zebrafish and potentially other fish impacted by columnaris disease.

**Figure 6.**
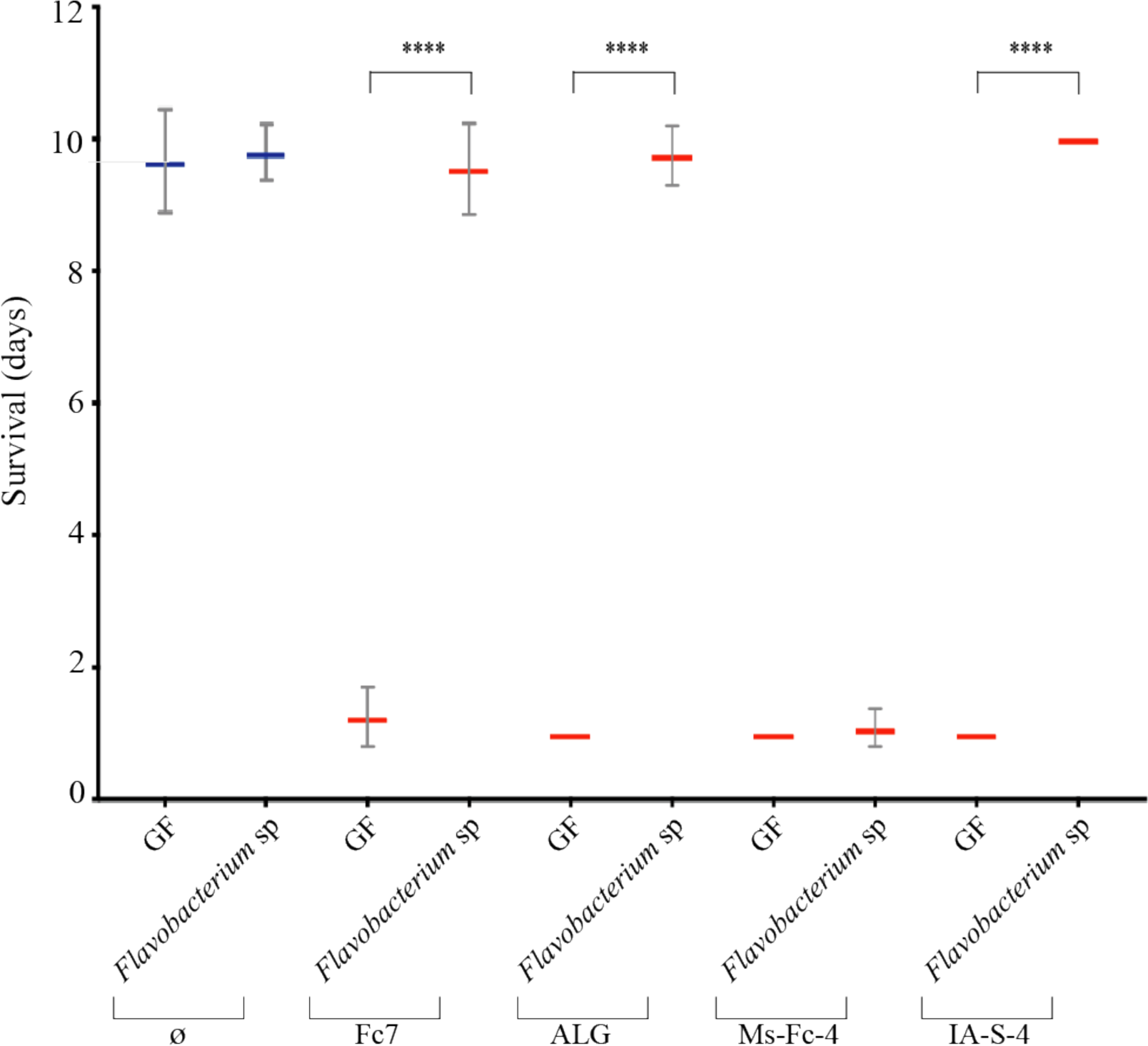
*Flavobacterium* sp. strain 4466 provides full protection to gnotobiotic zebrafish larvae against infection by three strains of *F. columnare*. Survival of GF zebrafish larvae exposed to *Flavobacterium* sp. strain 4466 48 h before infection with *F. columnare* strains Fc7, IA-S-4, Ms-Fc-4 and ALG-00-530. All *F. columnare* strains rapidly killed GF fish, whereas only strain Ms-Fc-4 rapidly killed fish that had been re-conventionalized with *Flavobacterium* sp strain 4466. Mean and SD plot representing average day post-infection at which infected fish died. For each condition n = 10 larvae. All surviving fish were euthanized at day 10. Asterisks indicate significant difference from non-infected population (****p<0.0001).

### Use of germ-free trout model to validate exogenous probiotics protecting against *F. columnare* infection

To determine whether our GF trout model could be used as a controlled gnotobiotic approach to screen for trout probiotics, we pre-exposed 22 dph GF rainbow trout larvae to *Chryseobacterium massiliae*, a bacterium that does not belong to trout microbiota but was previously shown to protect larval stage and adult zebrafish from infection by *F. columnare* [35]. After 48 h of bath in a *C. massiliae* suspension at 10^5^ CFU/ml, we infected trout larvae with *F. columnare* strains Fc7, ALG-00-530, IA-S-4 and Ms-Fc-4 and observed that *C. massiliae* protected against all tested *F. columnare* pathogens (Figure 7). These results showed that the GF rainbow trout model enables the evaluation of bacterial species, endogenous to trout or not, with probiotic potential against highly virulent *F. columnare* strains.

**Figure 7.**
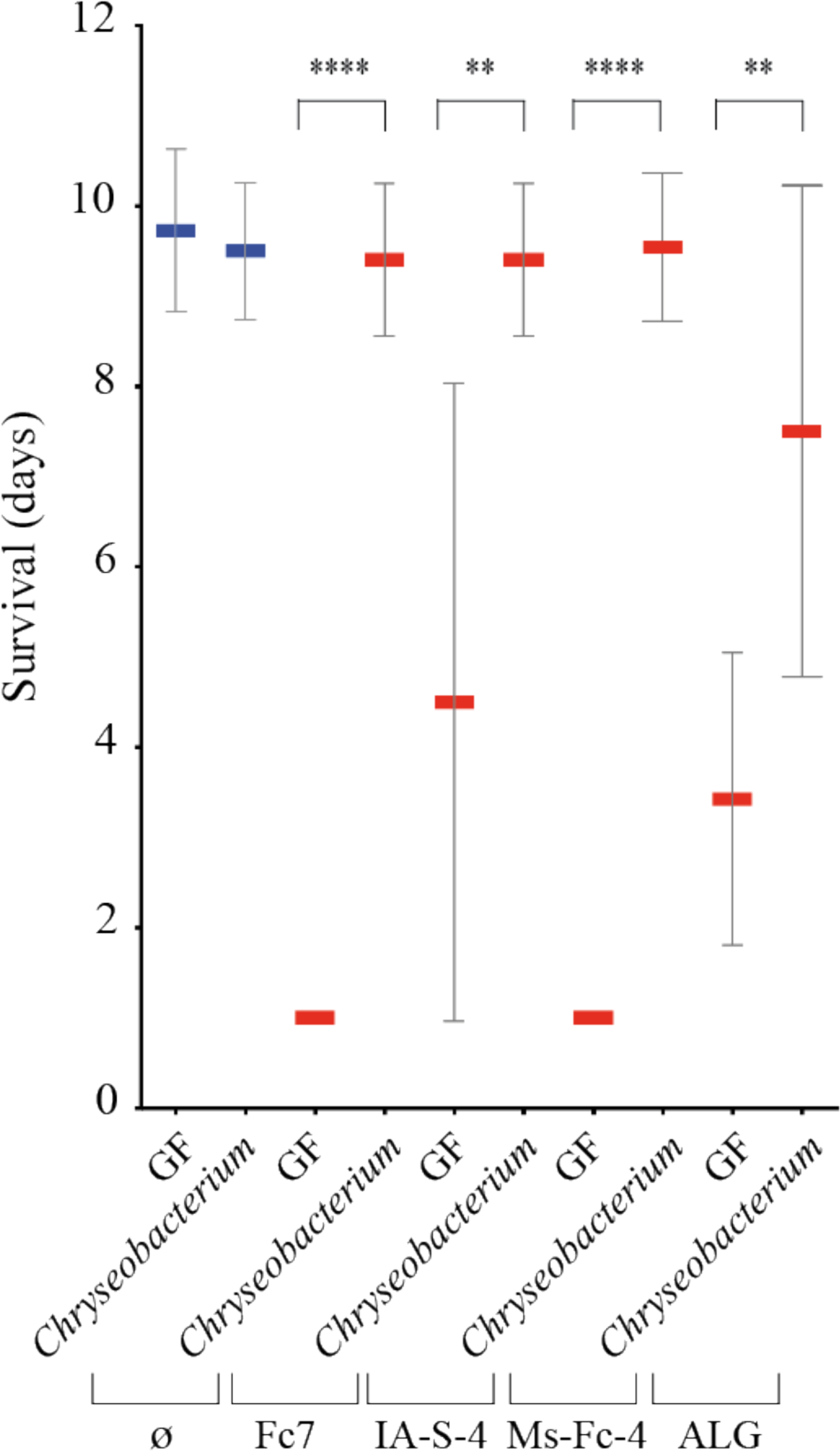
*C. massiliae* provides protection against *F. columnare* infection. GF larvae survival exposed to *C. massiliae* 48 h before infection with *F. columnare* strains Fc7, IA-S-4, Ms-Fc-4 and ALG-00-530. Mean and SD plot representing average day post-infection at which infected fish die. For each condition n = 10 larvae. All surviving fish were euthanized at day 10. Asterisks indicate significant difference from non-infected population (****p<0.0001; **p<0.01).

## DISCUSSION

Although the use of probiotics is a promising approach to improve fish growth and reduce disease outbreaks while limiting chemical and antibiotic treatments [17, 36, 37], rational and evidence-based procedures for the identification of protective bacteria are limited. Here, we established a controlled and robust model to study trout resistance to infection by bacterial pathogens and to identify trout probiotics in microbiologically controlled conditions using GF and gnotobiotic rainbow trout.

Our gnotobiotic protocol is based on the survival of rainbow trout eggs to chemical sterilization eliminating the microbial community associated to the egg surface. Similarly to gnotobiotic protocols used for zebrafish [24, 38], cod [25] and stickleback (*Gasterosteus aculeatus*) [39], our approach produced larvae that were GF up to 35 dph at 16°C without continued exposure to antibiotics, therefore avoiding possible effects of prolonged antibiotic exposure on fish development [40]. Similarly to GF stickleback larvae at 14 dph [39], we observed no development or growth differences between GF and Conv trout larvae at 21 dph. In contrast, GF sea bass (*D. labrax* L.) larvae grew faster and had a more developed gut compared to conventionally raised larvae [41]. These discrepancies could come from the fact that, in our study and in the GF stickleback study, anatomical analyses were performed before first-feeding, whereas the GF sea bass were already fed when examined [41]. Indeed, trout larvae initially acquire nutrients by absorbing their endogenous yolk until the intestinal track is open from the mouth to the vent. We therefore cannot rule out that at later stages of development, when fish begin to rely on external feeding, differences between GF and Conv fish may occur, especially in the structure and size of organs or in body weight. However, the hurdles associated with long-term fish husbandry while keeping effective sterility control, *de facto* limits our approach to relatively short-term experiments on larvae with limited feeding time and low complexity microbiota.

While GF conditions cannot be compared to those prevailing in the wild or used in fish farming [25], our results showed that GF rainbow trout larvae are highly susceptible to *F. columnare*, the causative agent of columnaris disease affecting many aquaculture fish species [30, 42]. Interestingly, our GF rainbow trout larvae model also revealed the protective activity of *C. massiliae*, a potential probiotic bacterium isolated from Conv zebrafish [35], against various *F. columnare* strains from different fish host and geographical origins. These results demonstrate that GF rainbow trout is a robust animal model for the study of *F. columnare* pathogenicity and support *C. massiliae* as a potential probiotic to prevent columnaris diseases in teleost fish other than its original host.

Furthermore, we demonstrated that the relatively simple culturable bacteria isolated from microbiota harbored by Conv trout larvae effectively protect against *F. columnare*. Interestingly, different studies have demonstrated that highly diverse gut communities are more likely to protect the host from pathogens [43, 44]. This constitutes the base for the paradoxical negative health effect associated with the massive utilization of antibiotics in aquaculture: the reduction in microbial diversity facilitates colonization by opportunistic pathogens [45]. While this advocates for practices leading to enrichment of fish microbial communities to minimize pathogenic invasions in aquaculture [16], our results demonstrate that resistance to a bacterial pathogen can also be achieved by a single bacterial strain in a low complexity microbiota. Moreover, previous studies of resistance to infection provided by controlled bacterial consortia in gnotobiotic hosts often relied on community composition, rather than individual members of the microbiota [46-49]. We showed that the observed protection in larvae is mainly due to the presence of *Flavobacterium* sp. strain 4466. We cannot exclude, however, that at later developmental stages, the presence of other bacterial species may be needed for more efficient implantation or stability of protective members in the trout microbiota.

The molecular basis of *F. columnare* pathogenicity is poorly understood, but was recently shown to rely on the secretion of largely uncharacterized virulence factors and toxins by the *Flavobacterium* type IX secretion system (T9SS) [50]. The high genetic variability of *F. columnare* and its broad host range constitute an important limitation for the identification of effective probiotics against this widespread pathogen. Several probiotic candidates isolated from the host provided partial protection against *F. columnare* infection in other conventional fish species such as walleye (*Sander vitreous*) and brook char (*Salvelinus fontinalis*) [51, 52]. However, high variability in protection provided by probiotic strains against *F. columnare* was observed depending on the fish batch used, indicating a resistance directly dependent on the fish host genetics [51] or immunological status. Here we reduced this variability using GF and gnotobiotic trout larvae and demonstrated the ability of *Flavobacterium sp*. strain 4466 isolated from Conv trout larvae microbiota to protect against *F. columnare* infection. Furthermore, this bacterium, but not its supernatant, inhibits *F. columnare* growth *in vitro*, which suggests a direct interaction between *Flavobacterium* sp. strain 4466 and *F. columnare*. Intriguingly, *Flavobacterium* sp. strain 4466 encodes a complete subtype T6SS^iii^, a molecular mechanism that delivers antimicrobial effector proteins upon contact with target cells and is unique to the phylum *Bacteroidetes* [53]. The members of *Flavobacterium* genus are ubiquitous inhabitants of freshwater and marine fish microbiota and both commensal and pathogenic *Flavobacterium* often share the same ecological niche [54-56]. Whether the *Flavobacterium* sp. strain 4466 T6SS^iii^ contact-dependent killing system contributes to colonization resistance by inhibiting *F. columnare* Fc7 growth is currently under investigation. We cannot, however, exclude other mechanisms such as competition for nutrients or pathogen exclusion upon direct competition for adhesion to host tissues. This process has been suggested for infected zebrafish with efficient colonization of highly adhesive probiotic strains and enhanced life expectancy [24, 57, 58].

For the past 30 years, the fish farming industry has dedicated considerable efforts to identify probiotic microorganisms for rainbow trout, including Gram-positive and Gram-negative bacteria and yeast [59]. However, the high interindividual and seasonal variability of trout microbiota [60, 61] and the random or time-limited colonization ability of exogenous microorganism rarely enables consistent probiotic efficacy. Despite some studies of rainbow trout proposing different endogenous bacterial strains as probiotic candidates, few have demonstrated protective properties against pathogenic bacteria *in vivo* [62-65]. Short-residing probiotics may limit unintended consequences to the microbial community and host system, but the use of endogenous residents may stably modulate the community and protect the fish against reoccurring disease outbreaks over longer timescales [66, 67]. The probiotic efficacy of *Flavobacterium* sp. strain 4466 against different strains of *F. columnare* from different fish hosts and geographical origins, suggests that it could be used as a broad probiotic to prevent infections.

In conclusion, we showed that germ-free and gnotobiotic trout larvae are an effective experimental tool to study microbiota-determined sensitivity to major salmonid freshwater pathogens, enabling the validation of endogenous and exogenous potential probiotic strains. This approach will also be instrumental in studying the molecular basis of probiosis against fish pathogens as well as host-pathogen mechanisms, ultimately contributing to the mitigation of rainbow trout diseases in aquaculture.

## MATERIAL AND METHODS

### Ethics statement

All animal experiments described in the present study were conducted at the Institut Pasteur according to European Union guidelines for handling of laboratory animals (http://ec.europa.eu/environment/chemicals/lab_animals/home_en.htm) and were approved by the Institut Pasteur institutional Animal Health and Care Committees under permit # dap200024

### Handling of rainbow trout larvae

Rainbow trout (AQUALANDE breeding line) “eyed” eggs of 210 to 230 degree-days (21-23 days after fertilization at 10°C) (dd) were obtained from Aqualande Group trout facility in Pissos, France. Upon arrival, the eggs were progressively acclimatized to 16°C before manipulation. All procedures were performed under a laminar microbiological cabinet and with single-use disposable plastic ware. Eggs were kept in 145 × 20 mm Petri dishes with 75 mL autoclaved dechlorinated water until hatching. After hatching, fish were transferred and kept in 250 mL vented cap culture flasks in 100 mL sterile water at 16°C. Fish were fed starting 21 days post-hatching with gamma-ray sterilized fish food powder every 48 h, 30 minutes before water renewal of half the volume of water to avoid waste (NH_4_^+^, NO_2_-, NO_3_-) accumulation and oxygen limitation.

### Sterilization and raising of germ-free rainbow trout

The eyed rainbow trout eggs received at 210 dd were transferred to sterile Petri dishes (140 mm diameter, 150 eggs/dish) and washed twice with a sterile methylene blue solution (0.05 mg/ml). The eggs, kept in 75 ml of methylene blue solution, were then exposed to a previously described antibiotic cocktail [24] (750 µl penicillin G (10,000 U/ml), streptomycin (10 mg/ml); 300 µl of filtered kanamycin sulfate (100 mg/ml) and 75 µl of the antifungal drug amphotericin B solution (250 µg/ml)) for 24 hours by gentle agitation at 16°C. Eggs were then washed 3 times with fresh sterile water and treated with bleach (0.005 %) for 15 minutes. Following 3 washes with sterile water, eggs were treated for 10 minutes with 10 ppm Romeiod (COFA, France), a iodophor disinfection solution. Finally, eggs were washed 3 times and kept in a class II hood at 16°C in 75 ml of sterile water supplemented with the previously mentioned antibiotic cocktail until hatching spontaneously 5 to 7 days following the disinfection process. Once hatched, fish were immediately transferred to 75 cm^3^ vented cap culture flasks containing 100 ml of fresh sterile water without antibiotics (12 larvae/flask). The hatching percentage was determined by comparing the number of hatched larvae in Petri dish relative to the total number of eggs. *Sterility*: Sterility was monitored by culture-based and 16S rRNA PCR-based tests at 24 h, 7- and 21-day post-treatment. After feeding started, 50 µl of GF fish flask water was sampled before each water change as well as one larva every week to perform culture-based and 16S rRNA-based PCR sterility tests. 50 µl of rearing water from each flask was plated on LB, YPD and TYES agar plates, all incubated at 16°C under aerobic conditions. Fish larvae were also checked for bacterial contamination every week using the following methods. Randomly chosen fish were sacrificed by an overdose of filtered tricaine methane sulfonate solution (tricaine, Sigma, 300 mg/L). Whole fish were mechanically disrupted in Lysing Matrix tubes containing 1 ml of sterile water and 425-600 µm glass beads (Sigma). Samples were homogenized at 6.0 m s^-1^ for 45 s on a FastPrep Cell Disrupter (BIO101/FP120 QBioGene) and serial dilutions of the homogenized solution were plated on LB, YPD and TYES agars. When water samples or fish homogenates showed any bacterial CFU on the different culture media used, the corresponding animals (or flasks) were removed from the experiment. The absence of any contamination in the fish larvae was further confirmed by PCR as follows. Total bacterial DNA was extracted from fish homogenate sample using QIAmp DNA Microbiome Kit (Qiagen) following manufacturer instructions. All reagents used were molecular grade and supplied by Sigma-Aldrich (UK). To detect the presence of microbial DNA, universal specific primers for the chromosomal 16S rRNA (27F: 5’-AGAGTTTGATCCTGGCTCAG-3’; 1492R 5’-GGTTACCTTGTTACGACTT-3’) were used for the PCR [68].

### Bacterial strains and growth conditions

*F. columnare* strains Fc7 [69], Ms-Fc-4 [70] and IA-S-4 [71] (genomovar I), ALG-00-530 [72] (genomovar II), and *Chryseobacterium massiliae* [35] were grown in tryptone yeast extract salts (TYES) broth [0.4% (w/v) tryptone, 0.04% yeast extract, 0.05% (w/v) MgSO_4_ 7H_2_O, 0.02% (w/v), CaCl_2_ 2H_2_O, 0.05% (w/v) D-glucose, pH 7.2] at 150 rpm and 18°C. *F. psychrophilum* strains THCO2-90 was grown in TYES broth at 150 rpm and 18°C. *Yersinia ruckeri* strain JIP 27/88 was grown in Luria-Bertani (LB) medium at 150 rpm and 28°C. *V. anguillarum* strain 1669 was grown in tryptic soy broth (TSB) at 150 rpm and 28°C. *L. garvieae* JIP 28/99 was grown in brain heart infusion (BHI) broth at 150 rpm and 28°C. When required, 15 g/L of agar was added to the broth media to obtain the corresponding solid media. Stock cultures were preserved at -80°C in the respective broth media supplemented with 15% (vol/vol) glycerol.

### Fish infection challenge

Pathogenic bacteria were grown in suitable media at different temperatures until advanced stationary phase. Then, each culture was pelleted (10.000 rpm for 5 min) and washed once in sterile water. Bacteria were resuspended in sterile water and added to culture flasks at a final concentration of 10^7^ CFU/ml. After 24 hours of incubation with pathogenic bacteria at 16°C, fish were washed three times by water renewal. Bacterial concentrations were confirmed at the beginning and at the end of the immersion challenge by plating serial dilutions of water samples on specific medium for each pathogen. Ten to twelve larvae were used per condition and experiment and each experiment was repeated at least twice. Virulence was evaluated by daily monitoring of fish mortality up to 10 days post-infection.

### Characterization of culturable conventional rainbow trout microbiota

To identify the species constituting the cultivable microbiota of Conv trout larvae, 3 individuals were sacrificed with an overdose of tricaine at 35 dph, homogenized following the protocol described above and serial dilutions of the homogenates were plated on TYES, LB, R2A and TS agars. The plates were incubated a 16°C for 48 to 72 hours. All morphologically distinct colonies (based on form, size, color, texture, elevation and margin) were then isolated and conserved at -80°C in respective broth medium supplemented with 15 % (vol/vol) glycerol. In order to identify individual morphotypes, individual colonies were picked for each morphotype from each agar plates, vortexed in 200 µl DNA-free water and boiled for 20 min at 90°C. Five µl of this bacterial suspension was used as template for colony PCR to amplify the 16S rRNA gene with the universal primer pair 27f and 1492R. 16S rRNA gene PCR products were verified on 1% agarose gels, purified with the QIAquick^®^ PCR purification kit and two PCR products for each morphotype were sent for sequencing (Eurofins, Ebersberg, Germany). Individual 16S rRNA-gene sequences were compared with those available in the EzBioCloud database [73]. A whole genome-based bacterial species identification was performed for *Flavobacterium* sp. strain 4466 with the TrueBac ID system (v1.92, DB:20190603) (https://www.truebacid.com/) [74]. Species-level identification was performed based on the algorithmic cut-off set at 95% ANI or when the 16S rRNA gene sequence similarity was >99%.

### Whole genome sequencing

Chromosomal DNA of *Flavobacterium* sp. strain 4466 isolated from rainbow trout larvae microbiota was extracted using the DNeasy Blood & Tissue kit (QIAGEN) including RNase treatment. DNA quality and quantity was assessed on a NanoDrop ND-1000 spectrophotometer (Thermo Scientific). DNA sequencing libraries were made using the Nextera DNA Library Preparation Kit (Illumina Inc.) and library quality was checked using the High Sensitivity DNA LabChip Kit on the Bioanalyzer 2100 (Agilent Technologies). Sequencing clusters were generated using the MiSeq reagents kit v2 500 cycles (Illumina Inc.) according to manufacturer’s instructions. DNA was sequenced at the Mutualized Platform for Microbiology at Institut Pasteur by bidirectional sequencing, producing 2 × 150 bp paired-end (PE) reads. Reads were quality filtered, trimmed and adapters removed with fastq-mcf [75] and genomes assembled using SPAdes 3.9.0 [76].

### Phylogenomic analysis

The proteomes for the 15 closest *Flavobacterium* strains identified by the ANI analysis were retrieved from the NCBI RefSeq database. These sequences together with the *Flavobacterium* sp. strain UGB 4466 proteome were analyzed with Phylophlan (version 0.43, march 2020) [77]. This method uses the 400 most conserved proteins across the proteins and builds a Maximum likelihood phylogenetic tree using RAxML (version 8.2.8) [78]. Maximum likelihood tree was boostrapped with 1000 replicates.

### Germ free rainbow trout microbial re-conventionalization

Each isolated bacterial species was grown for 24 hours in suitable medium at 150 rpm and 20°C. Bacteria were then pelleted, washed twice in sterile water and diluted to a final concentration of 5.10^7^ CFU/ml. At 22 dph, GF rainbow trout were mono-re-conventionalized by adding 1 ml of each bacterial suspension per flask (5.10^5^ CFU/ml, final concentration). In the case of fish re-conventionalization with bacterial consortia, individual bacterial strains were washed, then mixed in the same aqueous suspension, each at a concentration of 5.10^7^ CFU/ml. The mixed bacterial suspension was then added to the flask containing GF rainbow trout as previously described. In all cases, fish re-conventionalization was performed for 48 h and the infection challenge with *F. columnare* was carried out immediately after water renewal.

### Histological examination

Histological sections were used to compare microscopical lesions between GF and Conv fish following infection with *F. columnare*. Sacrificed animals were fixed for 24 hours in Trump fixative (4 % methanol-free formaldehyde, 1 % glutaraldehyde in 0.1 M PBS, pH 7.2) [79]. Whole fixed animals were washed 3 times for 30 min and 12 hours in 0.1 M of phosphate buffer, and post-fixed for 2 hours with 2 % osmium tetroxide (Electron Microscopy Science, Hatfield, PA, USA) in 0.15 M of phosphate buffer. After washing in 0.1 M of phosphate buffer for 2 ×10 min and 2 × 10 min in distillated water, samples were dehydrated in a graded series of ethanol solutions (50 % ethanol in water × 10 min; 70 % ethanol 3 × 15 min; 90 % ethanol 3 × 20 min; and 100% ethanol 3 × 20 min). Final dehydration was performed by 100 % propylene oxide (PrOx, TermoFisher GmbH, Kandel, Germany) 3 × 20 min. Then, samples were incubated in PrOx/EPON epoxy resin (Sigma-Aldrich, St Louis, MO, USA) mixture in a 3:1 ratio for two hours with closed caps, 16 hours with open caps, and in 100% EPON for 24 hours at room temperature. Samples were replaced in new 100% EPON and incubated at 37°C for 48 hours and at 60°C for 48 hours for polymerization. Semi-thin sections (thickness 1µm) and ultra-thin sections (thickness 70 nm) were cut with a “Leica Ultracut UCT” ultramicrotome (Leica Microsysteme GmbH, Wien, Austria).

Semi-thin sections were stained with toluidine blue solution for 1 min at 60°C, washed with distilled water for 5 seconds, ethanol 100 % for 10 seconds and distilled water again for 20 seconds, dried at 60°C and embedded in Epon resin which was allowed to polymerize for 48 hours at 60°C. Light microscopy images of semi-thin EPON sections were prepared with Nikon Eclipse 80i microscope connected with Nikon DS-Vi1 camera driven by NIS-ELEMENTS D4.4 (Nikon) software.

### Whole fish clearing and 3D imaging

For a 3D imaging of cleared whole fish, fish were fixed with 4 % formaldehyde in phosphate-buffered saline (PBS) overnight at 4°C. Fixed samples were rinsed with PBS. To render tissue transparent, fish were first depigmented by pretreatment in SSC 0.5X twice during 1 hour at room temperature followed by an incubation in saline sodium citrate (SSC) 0.5X + KOH 0.5 % + H_2_O_2_ 3 % during 2 hours at room temperature. Depigmentation was stopped by incubation in PBS twice for 15 minutes. Fish were then post-fixed with 2 % formaldehyde in PBS for 2 hours at room temperature and then rinsed twice with PBS for 30 min. Depigmented fish were cleared with the iDISCO+ protocol [80]. Briefly, samples were progressively dehydrated in ascending methanol series (20, 40, 60 and 80 % in H_2_O, then twice in 100 % methanol) during 1 hour for each step. The dehydrated samples were bleached by incubation in methanol + 5 % H_2_O_2_ at 4°C overnight, followed by incubation in methanol 100 % twice for 1 hour. They were then successively incubated in 67 % dichloromethane + 33 % methanol for 3 hours, in dichloromethane 100 % for 1 hour and finally in dibenzylether until fish became completely transparent. Whole sample acquisition was performed on a light-sheet ultramicroscope (LaVision Biotec, Bielefeld, Germany) with a 2X objective using a 0.63X zoom factor. Autofluorescence was acquired by illuminating both sides of the sample with a 488 nm laser. Z-stacks were acquired with a 2 µm z-step.

### Statistical methods

Statistical analyses were performed using unpaired, non-parametric Mann-Whitney test for average survival analysis and the log rank (Mantel-Cox) test for Kaplan–Meier survival curves. Analyses were performed using Prism v8.2 (GraphPad Software). A cut-off of p-value of 5 % was used for all tests. * p<0.05; ** p<0.01; *** p<0.001, **** p<0.0001.

## ACKNOWLEDGEMENTS

We thank Rebecca Stevick, Jean-Pierre Levraud, Pierre Boudinot, Eric Duchaud, Mark McBride and Christophe Beloin for critical reading of the manuscript. We thank Laurent Debruyne and Jérémie Vieville from the Pissos Aqualande Trout breeding station. We are grateful to Jean-François Bernardet and Mark McBride for kindly providing us some of pathogenic microorganisms used in this study. We thank Rustem Uzkebov for his help in histology analyses performed in the context of a service provided by the IBiSA Microscopy facility, Tours University, France. iDISCO imaging was established and performed by Christelle Langevin and Maxence Fretaud (INRAE EMERG’IN IERP phenotyping platform) and light-sheet images were acquired at the Institut de la Vision.

## DATA AVAILIBLITY STATEMENT

The genome of *Flavobacterium* sp. 4466 was deposited at European Nucleotide Archive (ENA) databank under the accession n° ERS4574862.

## AUTHOR CONTRIBUTIONS

D.P.-P. and J.-M.G. designed the experiments. J.B.-B., D.R. and J.-M.G. contributed to the initial experiments. D.P.P., S.V.-F and B.A. performed the experiments. D.P.-P. and R.P.N performed genomic analysis. D.P.-P. and J.-M.G. analyzed the data and wrote the paper.

## SUPPORTING INFORMATION

### SUPPORTING TABLES

**Supporting Table S1.** *Flavobacterium* sp. strain 4466 taxonomic identification based on genomic similarities. The identification was based on whole genome Average Nucleotide Identity (ANI), and percentage of similarity with 16S rRNA and *recA* genes. Whole genome-based bacterial species identification was performed by the TrueBac ID system.

| Taxon | ANI (%) | ANI coverage (%) | 16S *rRNA* (%) | *recA* (%) |
| --- | --- | --- | --- | --- |
| *Flavobacterium spartansii* | 94,65 | 82,4 | 97,80 | 98,51 |
| *Flavobacterium tructae* | 94,62 | 83,9 | 97,80 | 98,11 |
| *Flavobacterium chilense* | 85,26 | 39,8 | 97,27 | 89,48 |

**Supporting Table S2.**
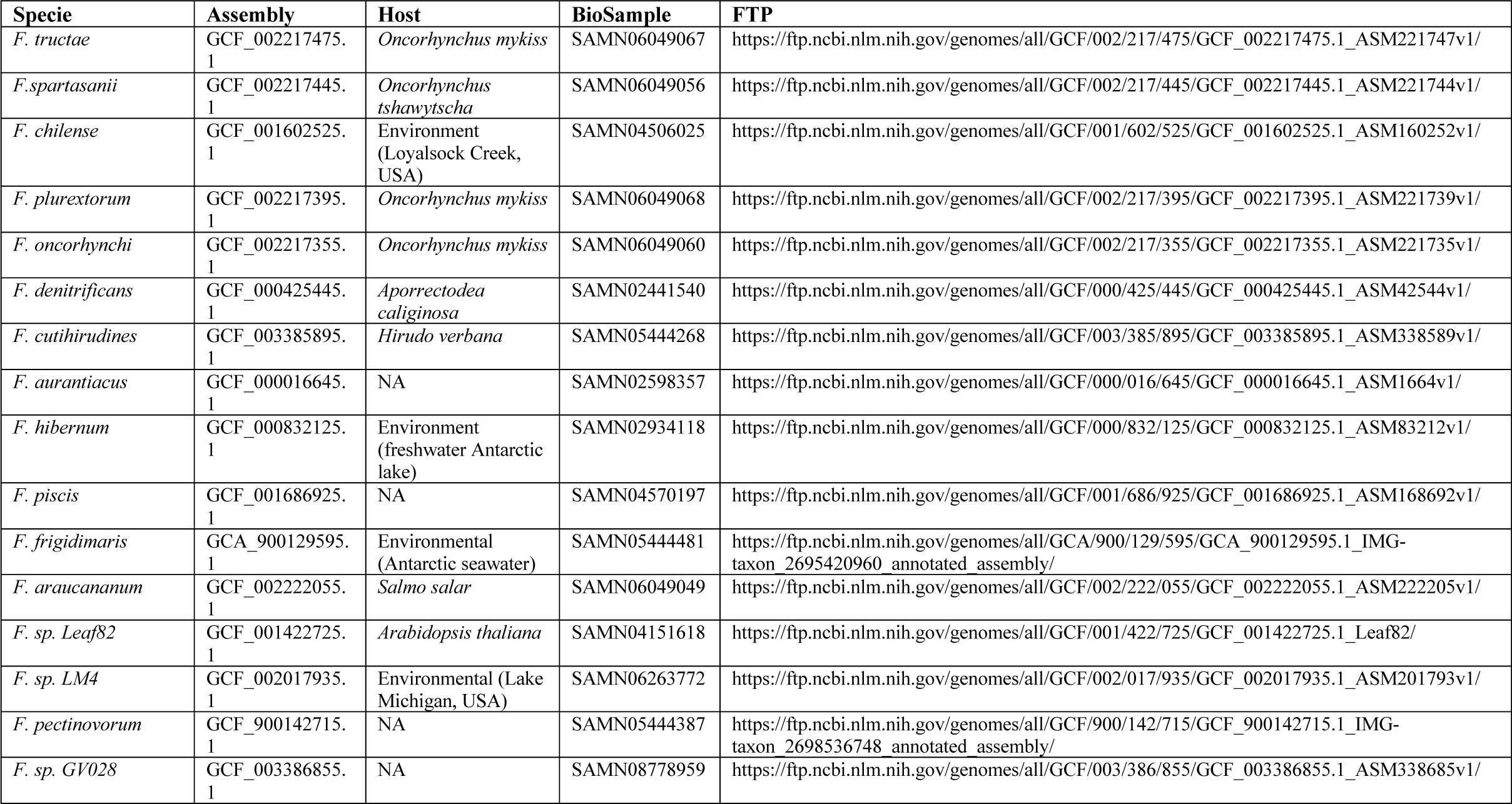
*Flavobacterium* species genomes retrieved from public databases.

### SUPPORTING FIGURES

**Supporting Figure S1.**
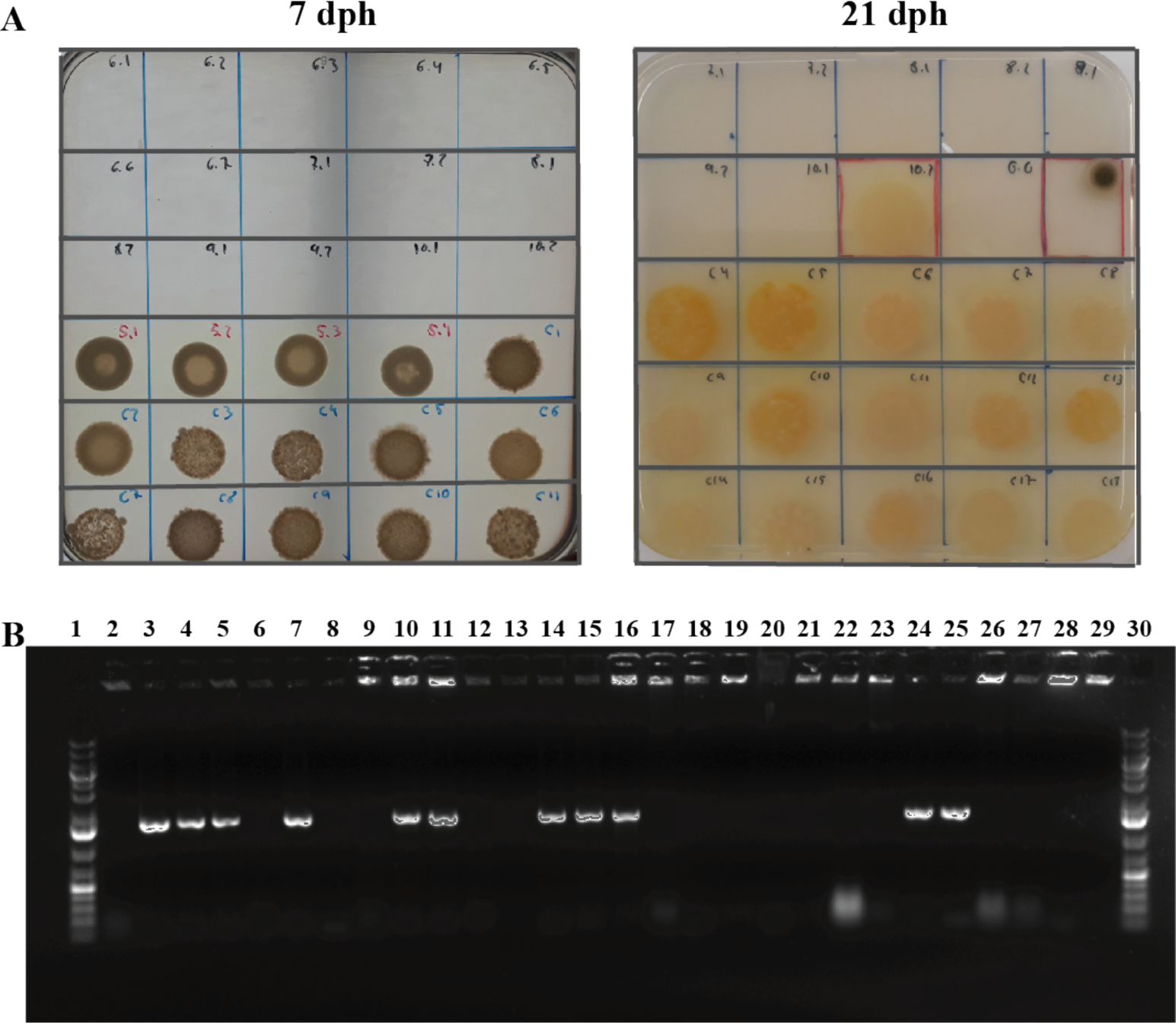
Sterility test of rainbow trout larvae raised under GF and Conv conditions. **A:** Culture-based sterility test of 50 µl samples of rearing water of GF and conventionally reared rainbow trout larvae at 7 and 21 dph. When water samples or fish homogenates showed bacterial CFU on any of the different culture media used, the corresponding animals (or flasks) were considered as non-sterile and removed from the experiment. **B**: PCR sterility test of total DNA extracted from 21 dph GF and conventionally reared rainbow trout larvae and used as a template for amplification of bacterial 16S rRNA gene. Lanes 1 and 30: molecular weight ladder; lane 2: non-template control; lanes 3-5: PCR products from Conv rainbow trout from three different flasks; lanes 6-29: PCR products from GF rainbow trout larvae from 23 different flasks. When water samples or fish homogenates showed a PCR amplification product, the corresponding animals (or flasks) were removed from the experiment.

**Supporting Figure S2.**
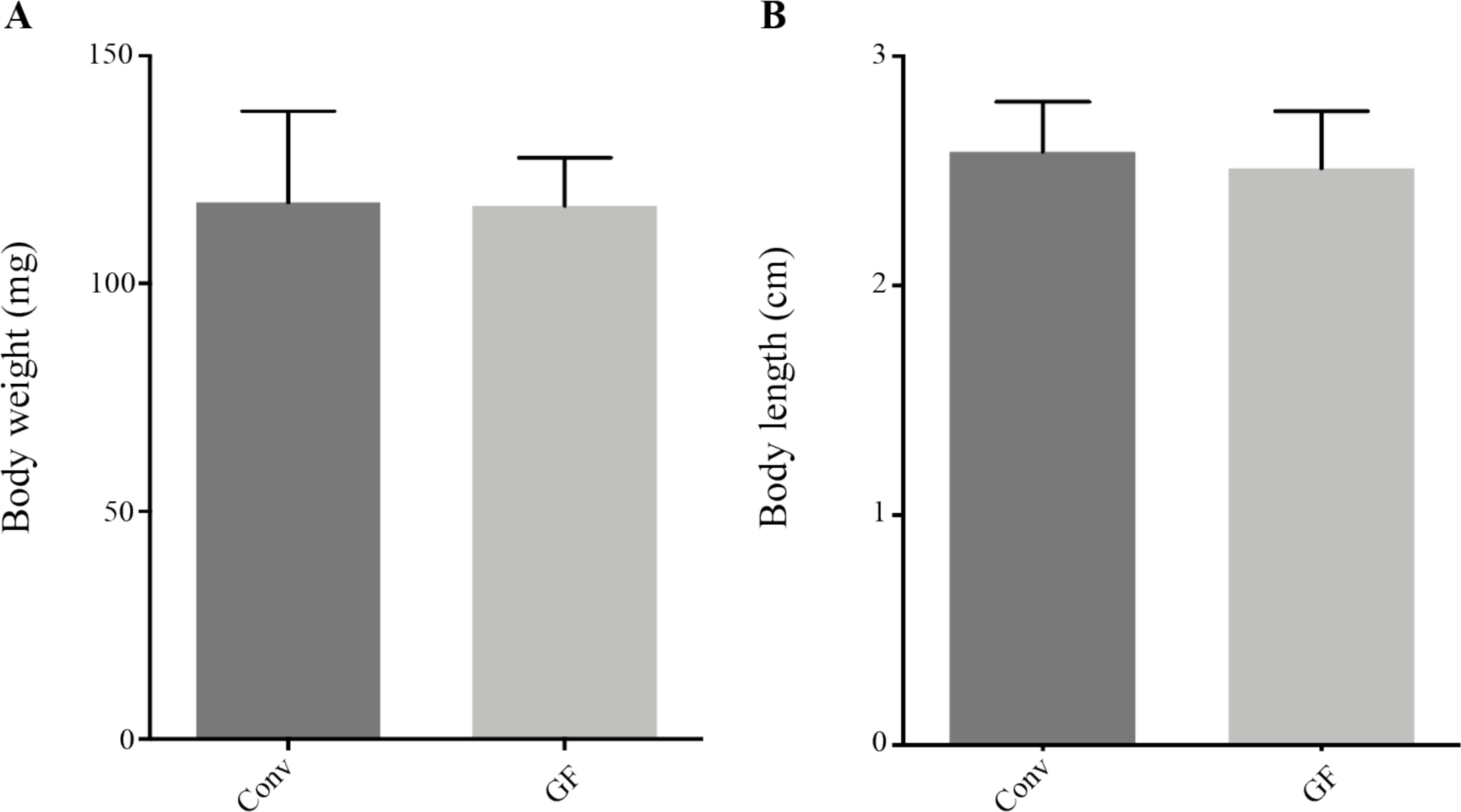
Growth performance of rainbow trout larvae raised under GF and Conv conditions. Conv and GF fish body size (A) and body weight (B) were measured at 35 dph (n= 5).

**Supporting Figure S3.**
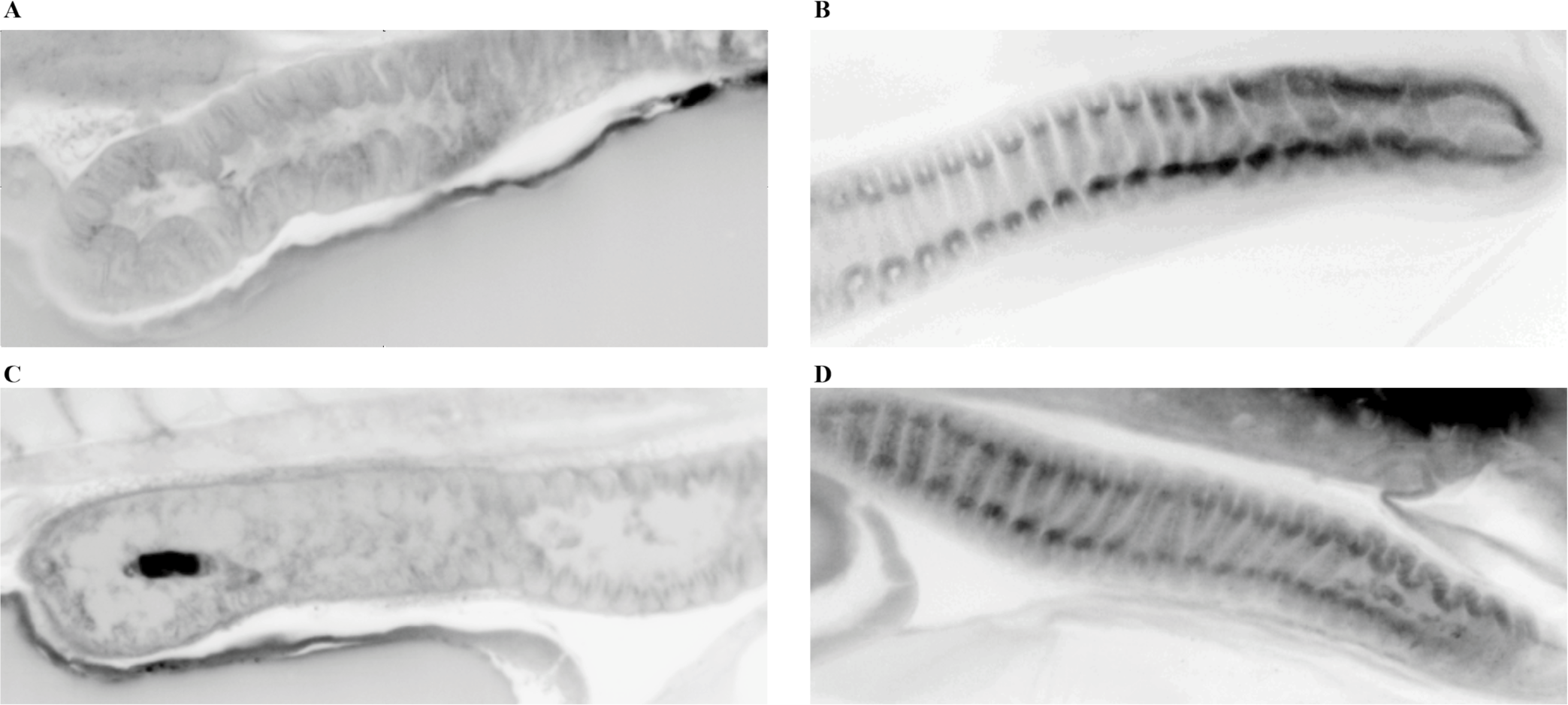
Anatomical comparison of the gut of Conv and GF rainbow trout larvae. 3D deep imaging of whole trout body corresponding to autofluorescence signal acquired by lightsheet microscopy after novel fish clearing processing. Selected optical sections of 21 dph gut were presented for Conv (A and B) and GF (C and D) rainbow trout larvae. Mid-gut (A and C), and posterior gut (B and D). Images representative of two different fish per condition.

**Supporting Figure S4.**
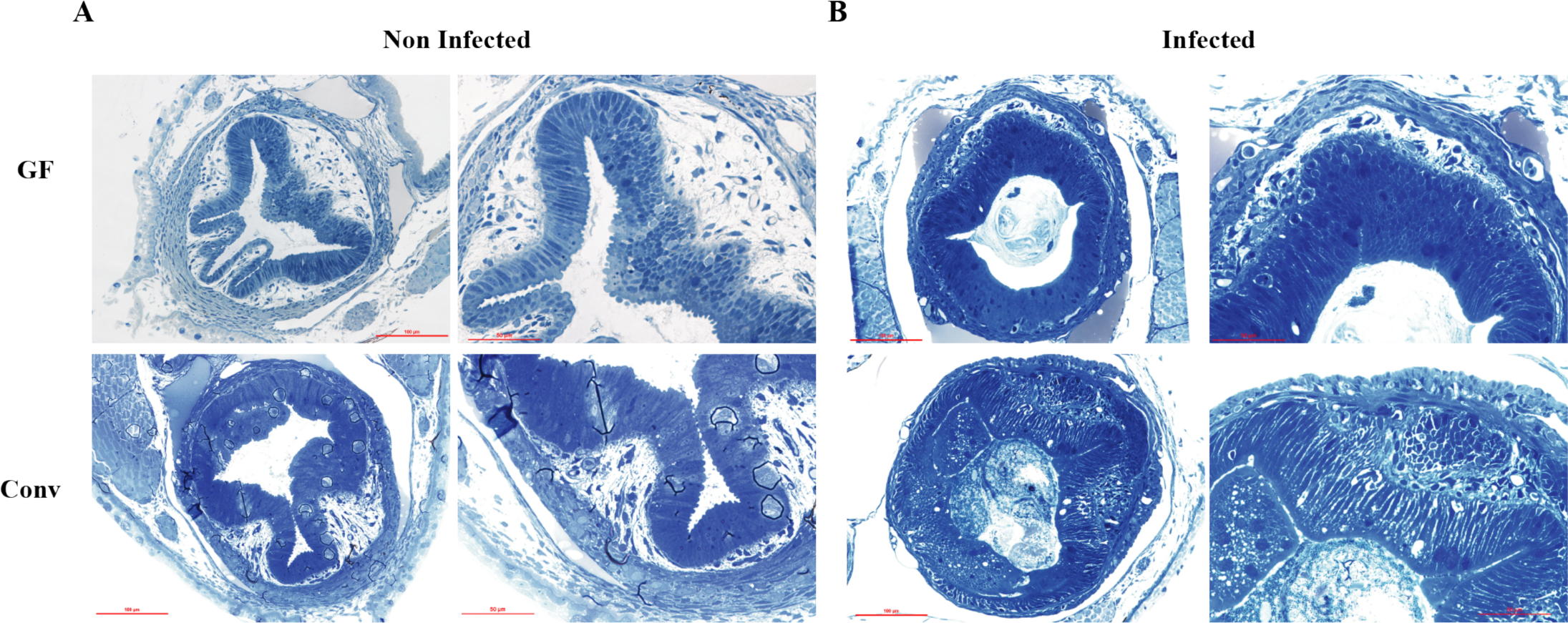
Histological comparison of the gut of infected and non-infected Conv and GF rainbow trout larvae. **A**: Representative images of intestines of non-infected larvae. **B**: Representative images of intestines of infected larvae exposed to *F. columnare* strain Fc7. Fish were fixed for histology analysis at 1 day post-infection (dpi). Toluidine blue staining of Epon-embedded zebrafish larvae for light microscopy.

**Supporting Figure S5.**
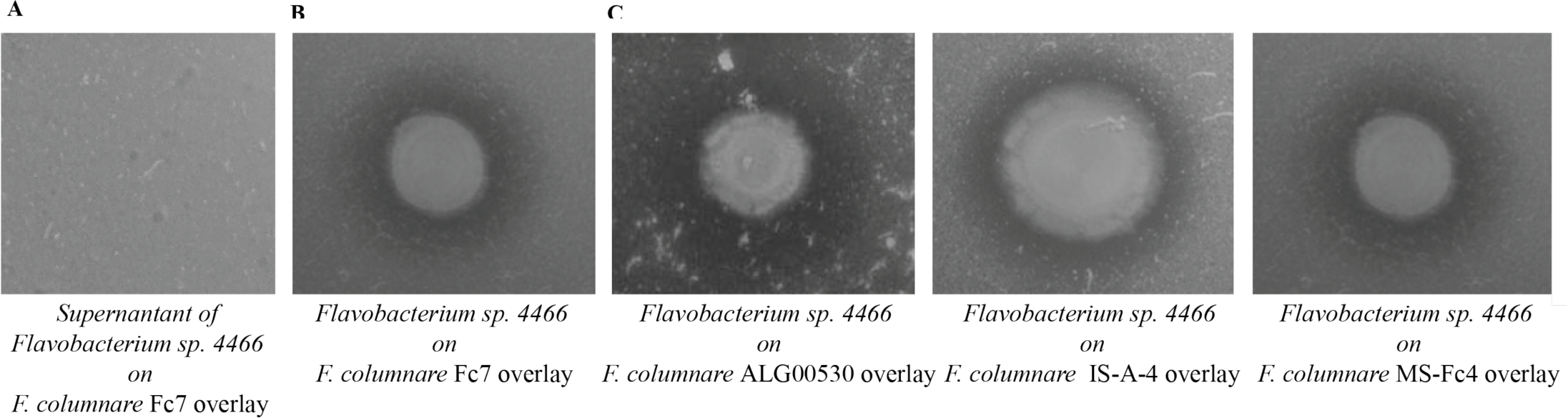
*In vitro* growth-inhibition activity of *Flavobacterium* sp. strain 4466 against different virulent *F. columnare* strains. **A:** lack of *F. columnare* Fc7 growth-inhibition after adding 5 µl of *Flavobacterium sp*. culture supernatant. **B:** Halo of *F. columnare* FC7 growth inhibition surrounding *Flavobacterium* sp. colony on a *F. columnare* strain Fc7 overlay. **C:** Halo of growth inhibition of *F. columnare* ALG-00-530, IA-S-4, and Ms-Fc-4. The agar overlay technique was performed by spreading *F. columnare* bacterial suspension on soft-agar solution over TYES agar, and then spotting 5 µl of an overnight culture of *Flavobacterium* sp. strain 4466. Incubation was performed at 28°C for 24 h.

**Supporting Figure S6.**
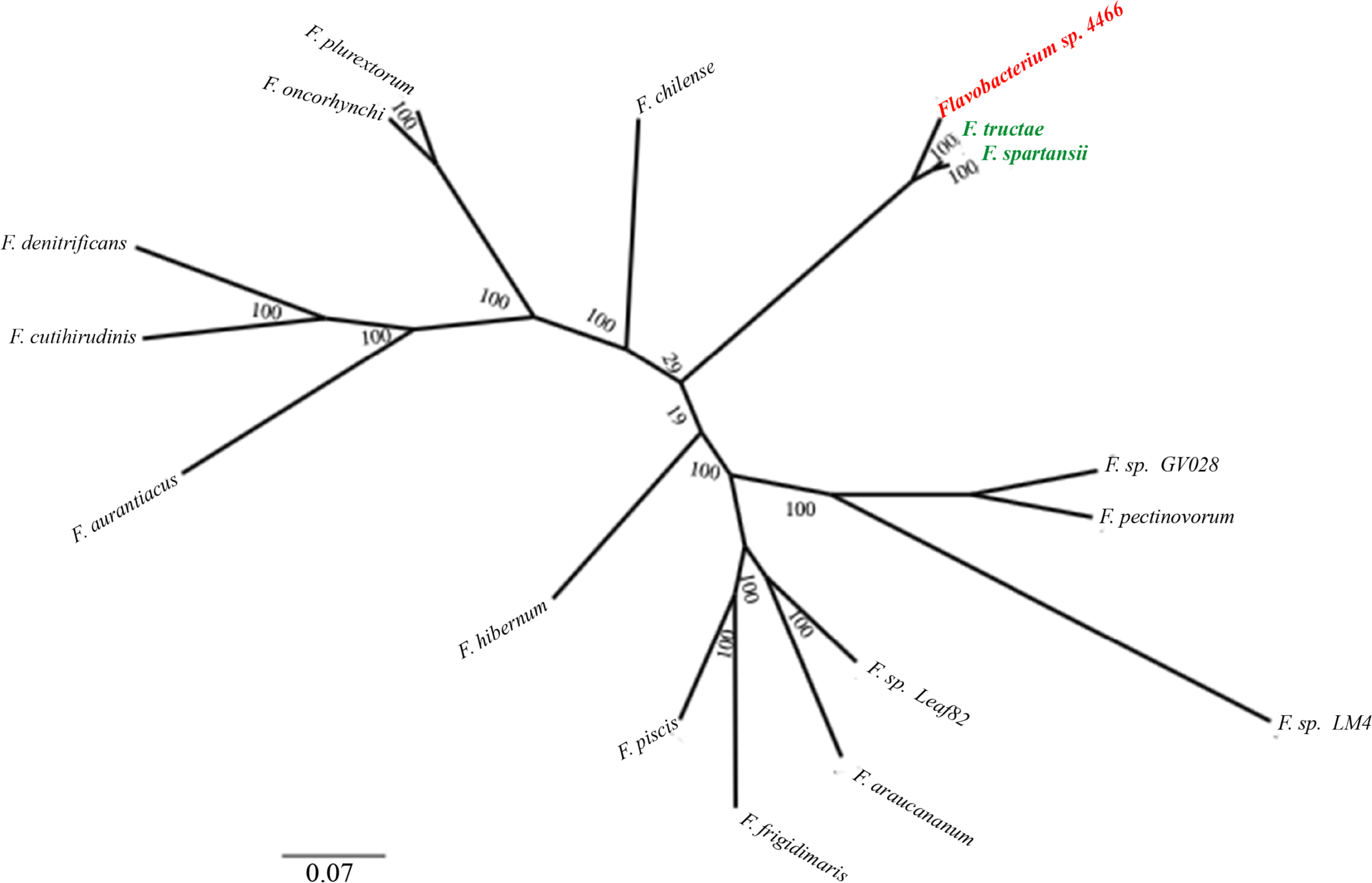
Phylogenetic tree illustrating the relationship between *Flavobacterium* sp. strain 4466 and the closest 15 *Flavobacterium* species based on ANI analysis. The tree was constructed with RAxML (version 8.2.8) by using the 400 most conserved proteins across the proteomes of each strain. Bootstrap support values are indicated in the nodes.

